# Multiple-Strain Infections of Human Cytomegalovirus with High Genomic Diversity are Common In Breast Milk from HIV-Positive Women in Zambia

**DOI:** 10.1101/493742

**Authors:** Nicolás M. Suárez, Kunda G. Musonda, Eric Escriva, Margaret Njenga, Anthony Agbueze, Salvatore Camiolo, Andrew J. Davison, Ursula A. Gompels

## Abstract

**Background:** In developed countries, human cytomegalovirus (HCMV) is a major pathogen in congenitally infected and immunocompromised individuals, in whom multiple-strain infection is linked to disease severity. The situation is less documente in developing countries. In Zambia, breast milk is a key route for transmitting HCMV and carries higher viral loads in HIV-positive women. We investigated HCMV strain diversity.

**Methods:** High-throughput sequence datasets were generated from 28 HCMV-positive breast milk samples donated by 22 mothers (15 HIV-positive and seven HIV-negative) at 4 or 16 weeks (or both) postpartum and analysed by genotyping 12 hypervariable HCMV genes.

**Results:** Among the 20 samples from 14 donors (13 HIV-positive and one HIV-negative) that yielded data meeting quality thresholds, 89 of the possible 109 genotypes were detected, and multiple-strain infections involving up to five strains per person were apparent in nine HIV-positive women. Strain diversity was extensive among individuals but conserved compartmentally and longitudinally within them. Genotypic linkage was maintained within the hypervariable UL73/UL74 and RL12/RL13/UL1 loci for virus-entry and immunomodulation, but not between genes more distant from each other.

**Conclusions:** Breast milk from HIV-positive women contains multiple HCMV strains of high genotypic complexity and thus constitutes a major source for transmitting viral diversity.

## BACKGROUND

Human cytomegalovirus (HCMV) is a major coinfection in HIV-positive people, in whom, as in other immunocompromised individuals such as transplant recipients, it contributes to morbidity and mortality. HCMV is also the most frequent congenital infection, causing adverse neurodevelopment, including hearing loss, microcephaly and neonatal morbidity. Postnatal infection generally occurs via milk in breastfeeding populations and is usually asymptomatic. However, it has been linked to morbidity, especially in preterm or underweight infants, and, in recent population studies, to adverse developmental effects, especially in association with HIV exposure in developing countries [1-4]. The most severe HCMV infections in transplant recipients, whether due to primary infection, reinfection or reactivation from latency, can result in severe or end-organ diseases such as retinitis, pneumonitis, hepatitis and enterocolitis [5]. Few studies of HCMV diversity, transmission and epidemiology have been conducted in relation to developing countries, including those having a high burden of endemic HIV.

HCMV has a double-stranded DNA genome of 236 kbp containing at least 170 protein-coding genes [6]. Diversity among strains is low overall, except in several hypervariable genes that exist as distinct, stable genotypes. These genes encode proteins that are particularly vulnerable to immune selection, including virus entry glycoproteins, other membrane glycoproteins and secreted proteins. The recombinant nature of HCMV strains was first identified in serological surveys and then in genomic studies, and is a key consideration for vaccine development [7-16]. However, understanding the pathogenic effects of HCMV diversity is at an early stage [17-19], and is limited by the fact that most analyses have focused on a few hypervariable genes characterised by polymerase chain reaction (PCR)-based genotyping [7, 12, 20]. This approach is relatively insensitive to the presence of minor strains in multiple-strain infections, which may have more serious outcomes.

High-throughput sequencing studies at the whole-genome level have started to facilitate a broader view of HCMV diversity, but most have involved isolating the virus in cell culture, which is prone to strain loss or mutation, or have depended on direct sequencing of PCR amplicons generated from clinical samples [11, 14, 16, 21]. Recent studies have avoided these limitations by using target enrichment to enable direct sequencing of strains present in clinical samples, most of which originated from patients in developed countries with congenital or transplantation-associated infections [11, 22-24]. We have used this approach to examine HCMV strain diversity in a developing country by analysing breast milk from women in Zambia, who constitute an HIV endemic population in sub-Saharan Africa, where we have previously demonstrated the negative developmental effects of early infection of infants with HCMV, particularly alongside HIV exposure [1, 3].

## METHODS

### Patients and samples

Anonymised breast milk samples were collected with informed consent as a substudy of the Breast Feeding and Postpartum Health study conducted at the University Teaching Hospital, Lusaka, Zambia, as approved by the ethical committees of the University Teaching Hospital and the London School of Hygiene and Tropical Medicine. This substudy included 28 HCMV-positive breast milk samples donated from one or both breasts by 15 HIV-positive and seven HIV-negative mothers at 4 or 16 weeks (or both) postpartum (**Table 1** and **Supplementary Table 1** rows 3-6) [3].

**Table 1.** Characteristics of donors, samples and datasets.

| Donor | HIV status | Breast sample | Weeks<br>postpartum | HCMV load<br>(ge/ml<sup>a</sup>) | Dataset |
| --- | --- | --- | --- | --- | --- |
| 158 | Negative | Left | 16 | 818,244 | 158L16 |
| 166 | Negative | Left | 16 | 282,252 | 166L16 |
| 193 | Negative | Right | 16 | 215,217 | 193R16<sup>b</sup> |
| 232 | Negative | Right | 4 | 470,150 | 232R4 |
| 239 | Negative | Right | 4 | 4,752,875 | 239R4 |
| 263 | Negative | Left | 4 | 5,285,775 | 263L4 |
| 280 | Negative | Right | 4 | 7,505,536 | 280R4 |
| 141 | Positive | Right | 16 | 319,888 | 141R16<sup>c</sup> |
| 154 | Positive | Left | 16 | 2,195,856 | 154L16 |
| 173 | Positive | Right | 16 | 349,532 | 173R16<sup>b</sup> |
| 174 | Positive | Left | 16 | 371,027 | 174L16<sup>b</sup> |
| 174 | Positive | Right | 16 | 642,516 | 174R16<sup>b</sup> |
| 181 | Positive | Left | 4 | 1,972,365 | 181L4 |
| 243 | Positive | Left | 16 | 643,895 | 243L16<sup>c</sup> |
| 243 | Positive | Right | 4 | 65,511,020 | 243R4<sup>b</sup> |
| 243 | Positive | Right | 16 | 795,092 | 243R16<sup>b</sup> |
| 248 | Positive | Right | 4 | 441,679 | 248R4<sup>b</sup> |
| 258 | Positive | Right | 4 | 610,613 | 258R4<sup>b</sup> |
| 259 | Positive | Left | 16 | 2,366,193 | 259L16<sup>b</sup> |
| 259 | Positive | Right | 16 | 5,053,047 | 259R16<sup>b</sup> |
| 264 | Positive | Left | 16 | 388,519 | 264L16<sup>b</sup> |
| 277 | Positive | Right | 16 | 250,377 | 277R16<sup>b</sup> |
| 278 | Positive | Left | 16 | 3,751,776 | 278L16<sup>b</sup> |
| 278 | Positive | Right | 4 | 294,246,272 | 278R4<sup>b</sup> |
| 278 | Positive | Right | 16 | 4,370,800 | 278R16<sup>b</sup> |
| 281 | Positive | Right | 4 | 20,291,530 | 281R4<sup>b</sup> |
| 283 | Positive | Right | 16 | 274,391 | 283R16<sup>b</sup> |
| 288 | Positive | Right | 4 | 31,574,022 | 288R4<sup>b</sup> |

<sup>a</sup>ge, genome equivalents.
<sup>b</sup>Met all quality thresholds.
<sup>c</sup>Met all quality thresholds except that unique fragment coverage depth was 10-20 rather than $\geq 20$ reads/nt.

### DNA extraction and viral load quantification

DNA was extracted from 200 µl breast milk using a QIAamp^®^ DNA mini kit (QIAGEN, Manchester, UK), and viral DNA load was measured using an in-house HCMV gB TaqMan assay run on an Applied Biosystems^®^ 7500 fast real-time PCR system (Applied Biosystems, Foster City, CA, USA), as described previously (**Table 1** and **Supplementary Table 1** row 7) [3].

### High-throughput DNA sequencing

The SureSelect^XT^ v. 1.7 target enrichment system (Agilent, Stockport, UK) was used to prepare sequencing libraries, as described previously (**Supplementary Table 1** rows 8-10) [22]. The libraries were sequenced using a MiSeq (Illumina, San Diego, CA, USA) with v. 3 chemistry to generate original datasets consisting of paired-end reads of 300 nucleotides (nt; **Table 1** and **Supplementary Table 1** rows 11-12).

### Phylogenetic analysis

UL73 and UL74 genotypes [7, 12] were investigated in 243 different HCMV strains [24]. MEGA 6.06 [25] was used to generate Muscle-derived amino acid sequence alignments and phylogenetic trees based on the Jones-Taylor-Thornton model and discrete gamma distribution with five categories.

### Strain characterisation using sequence motifs

Original datasets were quality-checked and trimmed using Trim Galore (http://www.bioinformatics.babraham.ac.uk/projects/trim_galore/; length=21, quality=10 and stringency=3) (**Supplementary Table 1** row 13). Bowtie2 [26] was used to remove reads mapping to the Genome Reference Consortium Human Reference 38 sequence, and also quality-checked and trimmed to create purged datasets (**Supplementary Table 1** row 14). Dataset quality parameters were (**Supplementary Table 1** rows 19-23) [24] on the basis of thresholds described in Results.

The number of genotypes was analysed by counting reads containing conserved, genotype-specific sequence motifs or their reverse complements. One short motif (14 nt) for each UL73 genotype and three short motifs (12-13 nt) for each UL74 genotype were identified by examining nucleotide sequence alignments and polymorphism plots derived from the 163 HCMV genome sequences in GenBank Release 211 (15 December 2015). Motif conservation was confirmed in the 243 genome set plus 383 UL73 and 72 UL74 single-gene sequences available in GenBank. The sequences of the short motifs are listed in **Table 2** with their frequency of occurrence.

**Table 2.**
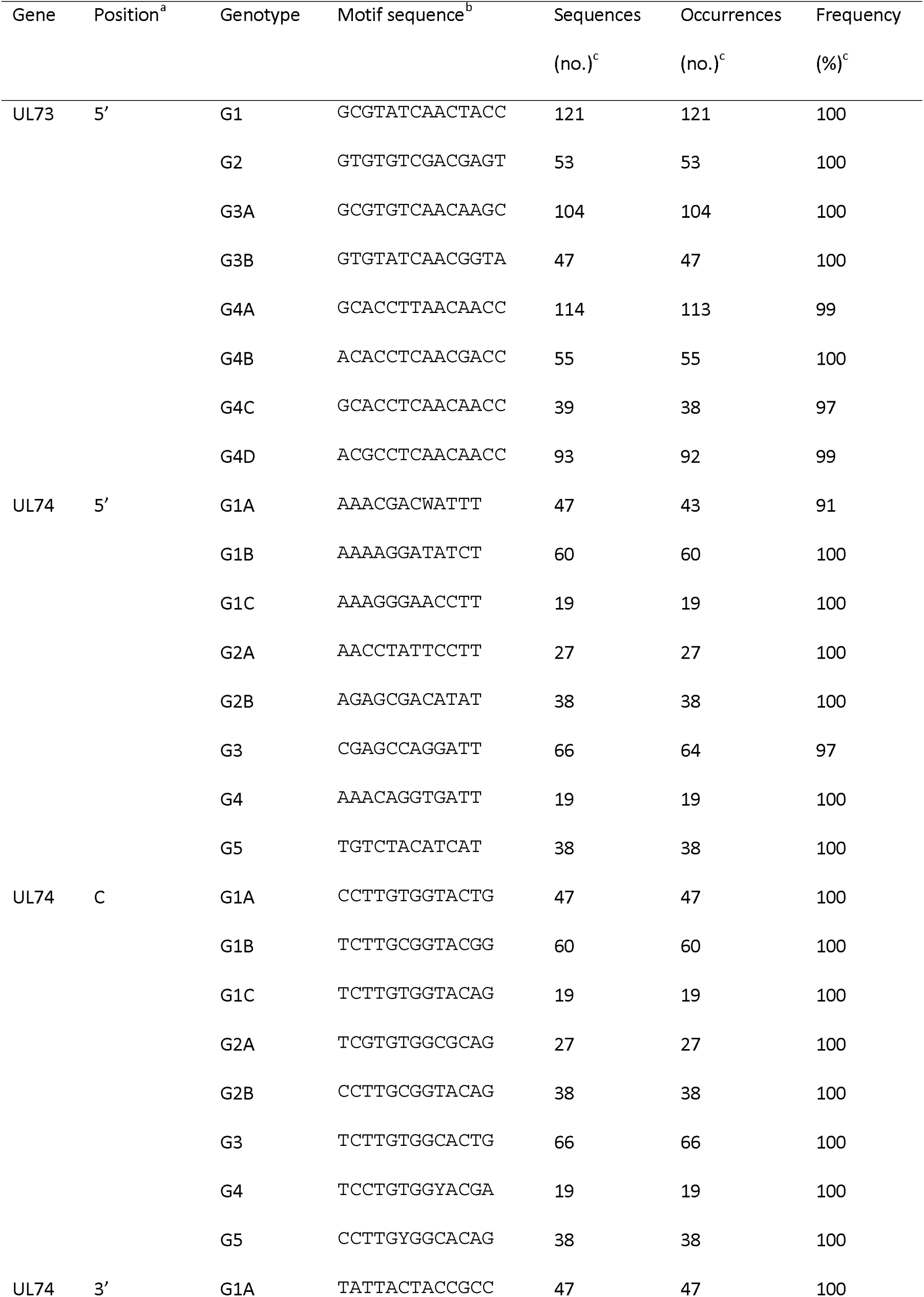
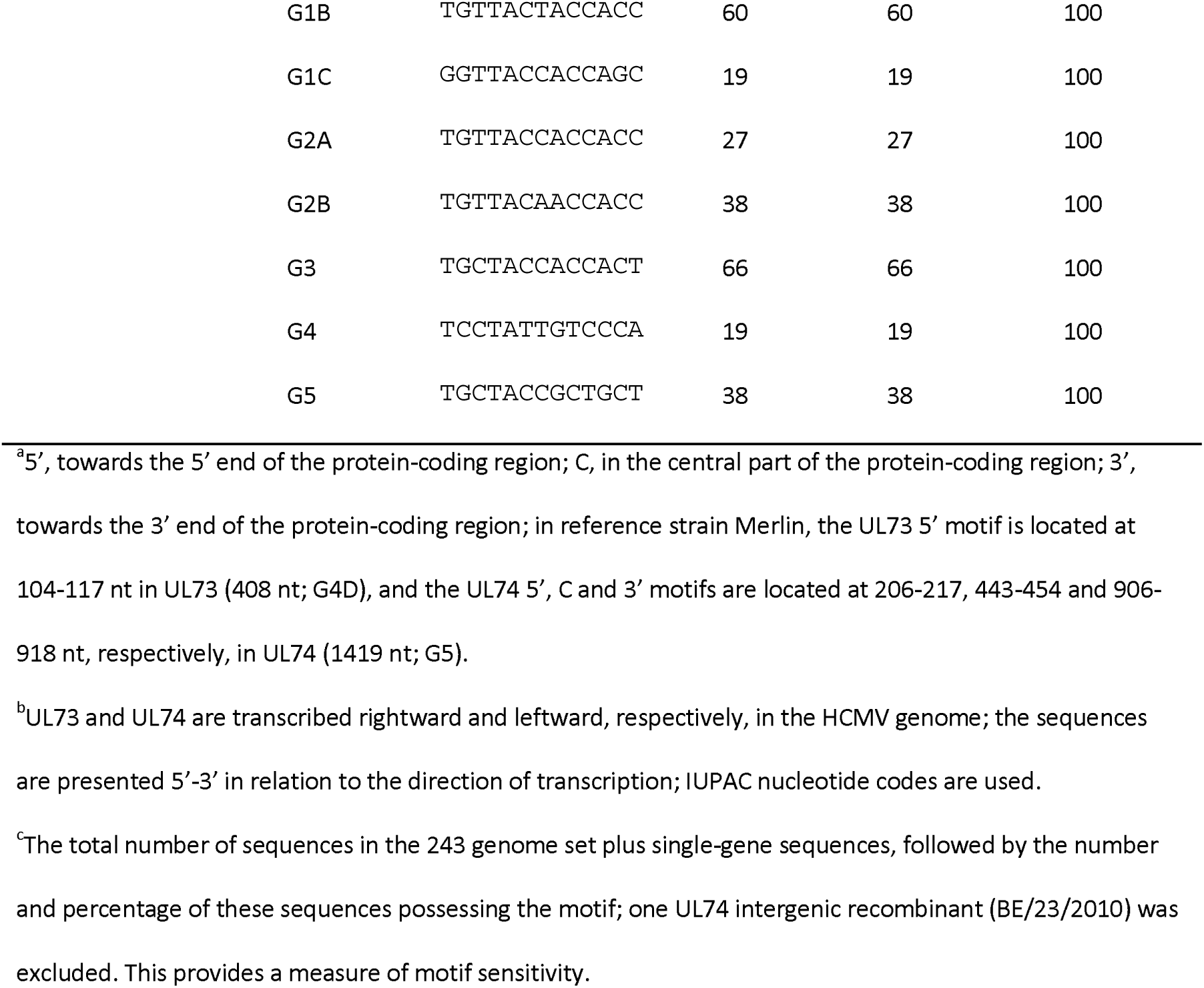
Short motif sequences in UL73 and UL74.

In addition to counting short motifs (**Supplementary Table 1** rows 25-56), long motifs (20-24 nt) at one per genotype were counted in UL73 and UL74 and a further ten hypervariable genes (RL5A, RL6, RL12, RL13, UL1, UL9, UL11, UL120, UL146 and UL139) (**Supplementary Table 1** rows 58-166). Long motifs identifying common mutations in three genes (RL5A, UL111A and US9) were also counted (**Supplementary Table 1** rows 167-174). The derivation of the long motifs as described [24], were used on the basis of thresholds described in Results (**Supplementary Table 1** row 17).

### Variant analysis

Replacement of one strain by another as the major population (genotype switch) in compartmental or longitudinal samples from the same individual were investigated by variant analysis [21, 27]. The original datasets were quality checked using FASTQC (http://www.bioinformatics.babraham.ac.uk/projects/fastqc/), trimmed to over 100 nt using Trimmomatic [28], optimised using VelvetOptimiser parameters (http://www.vicbioinformatics.com/software.velvetoptimiser.shtml), with assembly *de novo* using Velvet [29] and produced contigs checked by reference genome mapping with ABACAS [30]. The resulting contigs were verified by reference mapping using BWA [31] and SAMtools/BCFtools [32]. GATK [33] was used for indexing, mapping and variant calling, defining variant nucleotides as follows: prevalence <50%, overall read depth ≥50, average nucleotide quality ≥30, variant frequency ≥1% for read depths >1000 and >10% for read depths 50-1000, and minimum SNP depth ≥10. Artemis [34] was used for visualisation.

### Data deposition

The human purged datasets were deposited in the European Nucleotide Archive (ENA) under project number PRJEB31143 (**Supplementary Table 1** row 15). Complete genome sequences were assembled as described [22] and deposited in GenBank (**Supplementary Table 1** row 16).

## RESULTS

### Assessment of the sequence datasets

In a recent study, we highlighted the importance of monitoring the quality of datasets produced directly from clinical material by target enrichment and high-throughput sequencing [24]. We implemented this here by assembling the datasets against the reference strain Merlin genome (GenBank accession AY446894), noting the number of matching HCMV reads (**Supplementary Table 1** line 19), and deriving two parameters: (1) the percentage of matching HCMV reads in the dataset and (2) the percentage of the reference genome represented (**Supplementary Table 1** lines 20-21). Also, since sequencing methodology is highly PCR-based, the number of HCMV fragments from which the data were produced was monitored by additional parameters: (3) the coverage depth of the reference genome by all HCMV reads, and (4) the coverage depth of the reference genome by reads generated from unique HCMV DNA fragments (**Supplementary Table 1** lines 22-23).

Quality threshold values were set at: (1) ≥50%, (2) ≥95%, (3) ≥1000 of the total fragment reads/nt, and (4) ≥20 unique fragment reads/nt. Eighteen datasets generated from 13 women met all four criteria, and two datasets (141R16 and 243L16) met criteria (1) to (3) but exhibited lower values (between 10-20) for criterion (4). These 20 datasets (one from an HIV-negative women and 19 from 13 HIV-positive women) are indicated in **Table 1** and **Supplementary Table 1** row 11, and further analysed.

### Genotypic structure of the UL73/UL74 locus

Our previous study involving Sanger sequencing of single HCMV genes in breast milk samples obtained at multiple time points postpartum pointed to the presence of multiple strains [3]. We extended this by using sequence differences between the genotypes of hypervariable genes across the genome to characterise the strains represented in the datasets. We focused first on UL73 and UL74, as our earlier work had shown that these adjacent genes are markedly hypervariable, are almost always genotypically linked, and group into eight genotypes also identified in milk samples [3, 7, 12]. The nucleotide sequences were extracted from the set of 243 genome sequences and analysed phylogenetically (Figure 1). This confirmed the existence of eight genotypes for each gene (**Table 2**), the strong linkage between them (only seven recombinants were noted), and the high levels of intergenotypic diversity and low levels of intragenotypic diversity observed initially in small datasets [7, 12]. In the UL73 phylogeny, a single G4B strain (HAN; GenBank accession KJ426589) fell outside the genotypes due to three nucleotide differences that are characteristic of G4A strains and probably represent homoplasies. In the UL74 phylogeny, a single strain (BE/23/2010; GenBank accession KP745697) fell outside the genotypes and may have resulted from intragenic recombination between G1C and G1A. The distances between genotypes and branching in the two phylogenies also supported our previous inference that an ancestral recombination event had given rise to the linkage between UL73 G4C and UL74 G1C [12].

**Figure 1.**
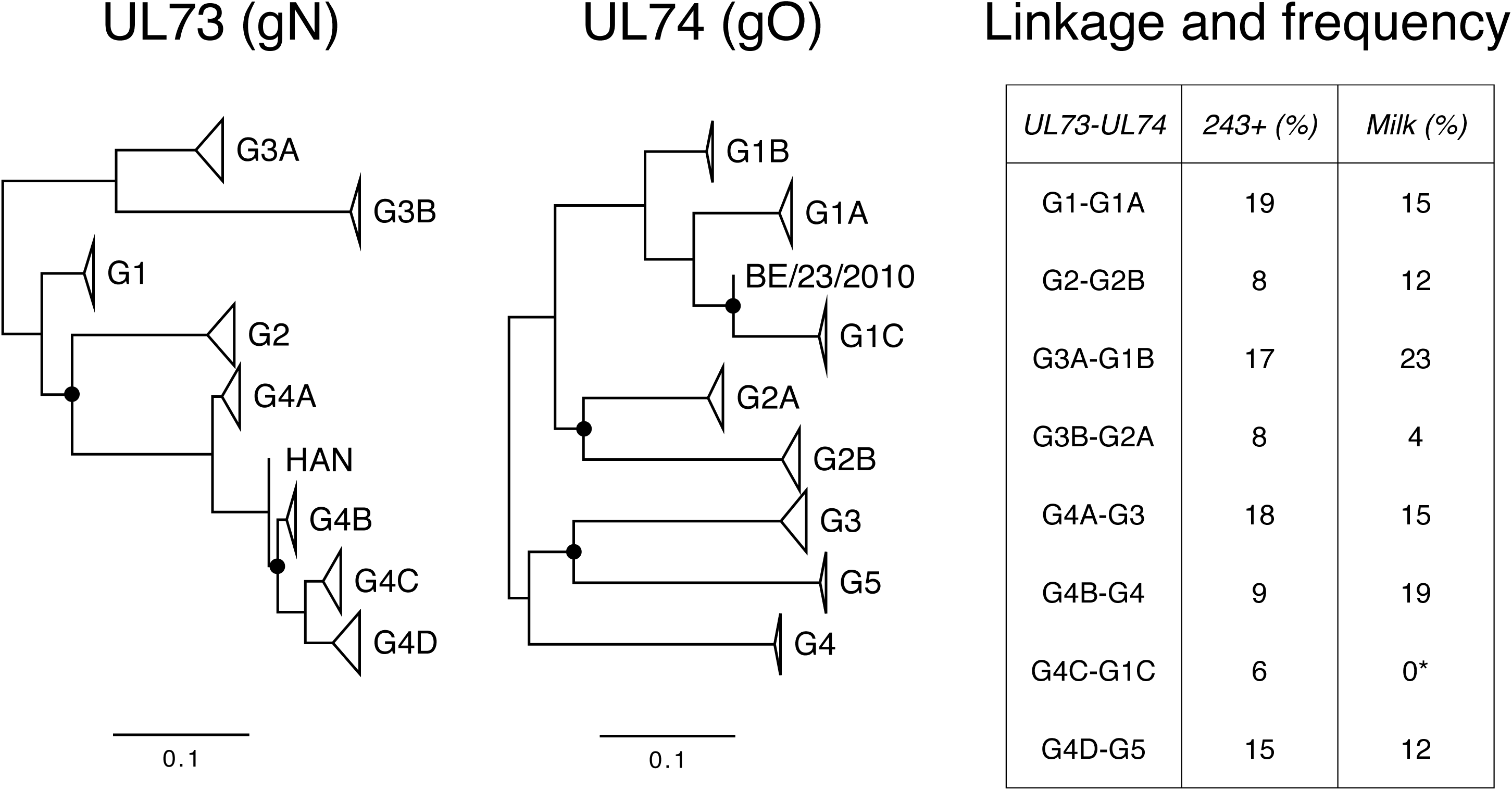
Unrooted phylogenetic trees for UL73 and UL74 based on amino acid sequences derived from 243 genome sequences, and a summary of genotypic linkages and frequencies. The site coverage cutoff value was 95%, leaving 134 sites in the UL73 tree (log likelihood-1117.87) and 435 sites in the UL74 tree (log likelihood-4153.86). Branch point robustness was inferred from 100 bootstrap replicates, and values of <70% are denoted by filled circles. Genotype branches are collapsed, and the numbers of substitutions per site are shown by the scale. The UL73 sequence of strain HAN and the UL74 sequence of strain BE/23/2010 did not fall into the genotypes. The linkages between UL73 and UL74 genotypes are listed, followed by the frequencies of UL73 genotypes in the 243 genome sequences plus single-gene sequences (243+; 626 in total; **Table 2**), and the frequencies of the deduced linkages in the samples (Milk; 26 in total; Table 3). The frequency of each genotype in the Milk set was not significantly different (*p*=0.05) from that in the 243+ set, as determined by random subsampling analysis (10,000 samplings of 26 genotypes from the set of 626), although there were some differences compared to the 243 genome set as noted in the text. Although no examples of this linkage were present in the datasets at levels in excess of the thresholds, at least one patient (258) was infected at sub-threshold levels by a relevant strain (**Supplementary Table 1**).

**Table 3.**
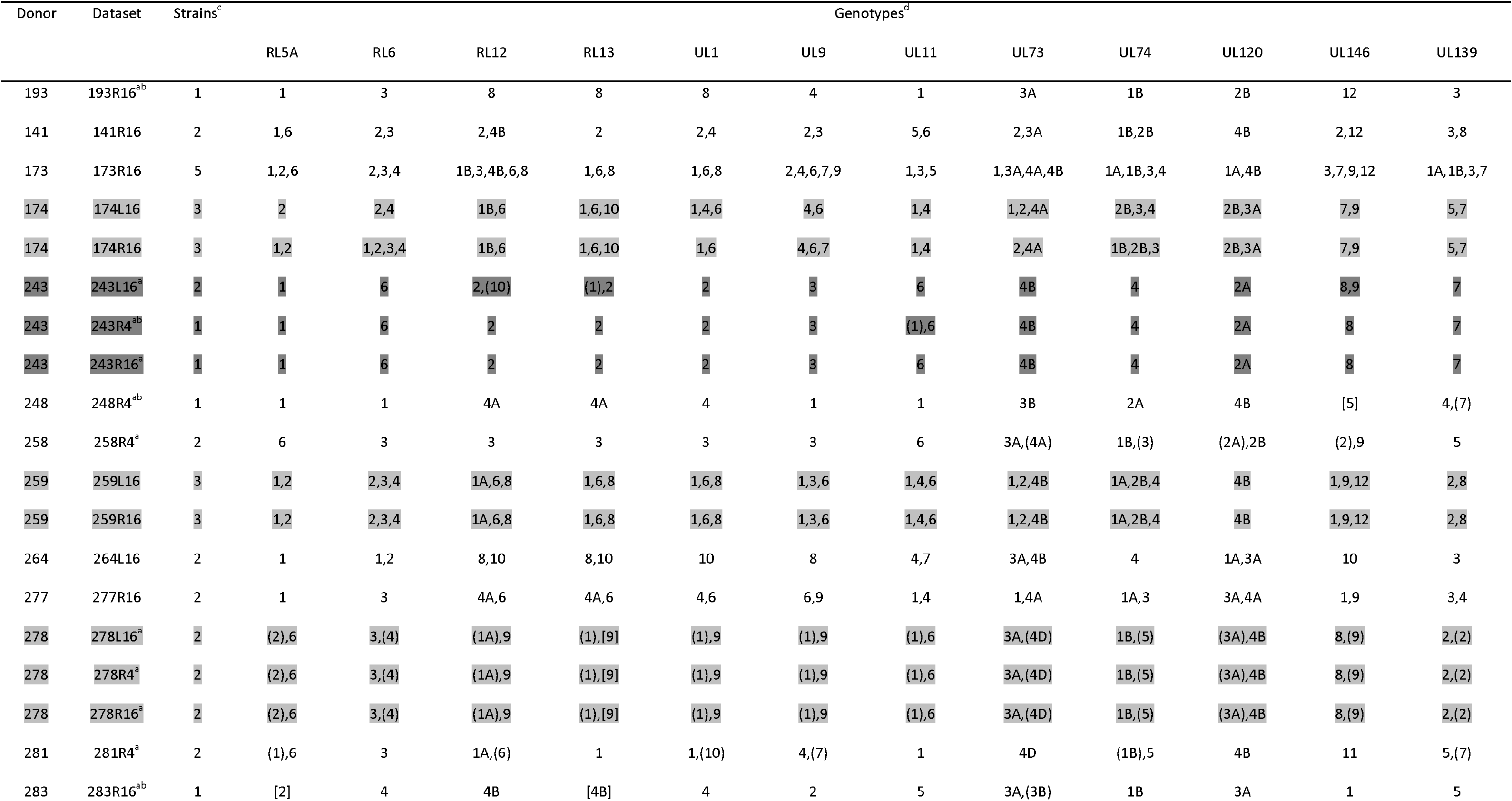
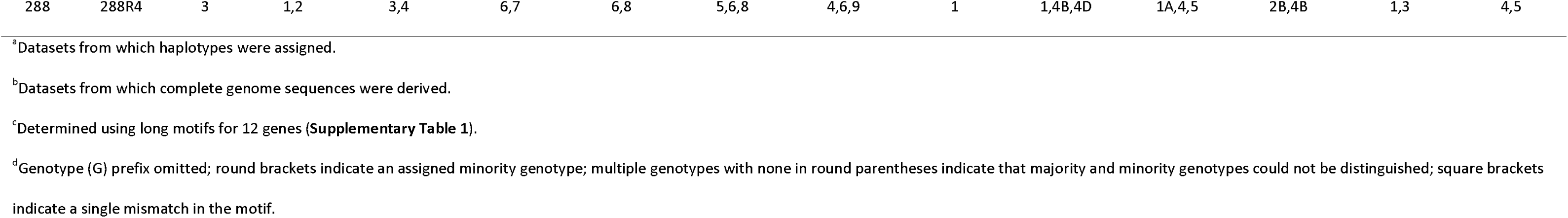
Genotypes and haplotypes assigned to datasets.

### Genotyping using sequence motifs

Having established a comprehensive view of UL73 and UL74 hypervariation, we developed short motifs capable of identifying individual genotypes. These consisted of a single motif near the 5’ end of each UL73 genotype and three separate motifs near the 5’ and 3’ ends and in the central region of each UL74 genotype. These motifs genotyped successfully the majority of the sequences used in the phylogenetic analyses (**Table 2**). We then extended the analysis to a further ten hypervariable genes, using a single, long, nonredundant motif for each genotype to improve discrimination.

The original datasets were trimmed (to create trimmed datasets) or purged of human reads and trimmed (to create purged datasets). The relative frequencies of individual genotypes were then estimated by counting motifs in all datasets with threshold requirements applied (**Supplementary Table 1** lines 25-56 and 58-166, respectively). Exclusion of human reads had little effect, except when short motifs were used with datasets containing a higher proportion of human reads. The UL74 5’ motif offered the least accurate genotypic discernment in such samples, perhaps as a result of its minimal length (12 nt). The number of strains in each sample was scored from the purged datasets using the long motifs with threshold requirements (**Table 3** and **Supplementary Table 1** row 17). A genotype was considered to be present when represented by >25 reads and >5% of the total number of reads detected for all genotypes of that gene, and the number of strains was scored as being the greatest number of genotypes detected using long motifs for at least two genes. Thus, strains present at <5% were unlikely to have been scored. There was a high degree of congruence between the results obtained using short and long motifs with datasets meeting the quality thresholds (**Supplementary Table 2**).

### Strain complexity in HIV-positive women

The majority of HIV-positive women (11/13) were infected by multiple HCMV strains (**Table 3** and **Supplementary Table 1**). The mode was at least two strains, and one woman was infected by five strains. The only HIV-negative woman was infected by a single strain, but it was not possible to discern from this whether multiple strains are more common in HIV-positive women. (The data from the other six HIV-negative women that were excluded from the analysis also indicate single infections in this group, but failure to meet the quality thresholds, partially from lower viral loads, could lead to underestimations). Even among this small cohort, 89 of the 109 possible genotypes of the 12 genes were detected. It was possible to assign fully linked genotypes (haplotypes) to eight strains represented in 11 datasets from seven donors, on the basis of complete genome sequences (four datasets) or the presence of a single strain or major and minor strains (when the former was highly predominant) in multiple-strain infections (**Table 3**). Consideration of other datasets indicated partially linked genotypes for up to 30 strains (**Supplementary Table 3**).

Genotypic linkage was detected only in two loci within which recombination has been shown to occur rarely, namely those containing the two respective sets of adjacent, hypervariable genes UL73/UL74 [12, 17, 35] and RL12/RL13/UL1 [11, 16]. The overall frequencies of UL73/UL74 genotypes in the milk samples were not significantly different from those in the 243 genome set plus single-gene sequences (Tables 1 and 3). There was some evidence for increased proportions of UL73/74 G4B-G4 and RL12/RL13/UL1 G2-G2-G2 in milk (p=.001 and p=0.02 respectively, compared to the 243 genome set), but case-controlled cohorts are required to confirm.

The use of three short motifs in UL74 facilitated an examination of intragenic recombination, and confirmed that strain BE/23/2010 is a recombinant with a G1C motif near the 5’ end and G1A motifs in the central region and near the 3’ end. In addition, compartmental stability was revealed by the use of both short and long motifs, in the form of genotypic conservation in samples from both breasts of four HIV-positive women (Figure 2). Small differences may be accounted for by minor strains present at levels nearing the detection threshold. Longitudinal stability was observed in two donors (243 and 278) with samples taken at weeks 4 and 16 postpartum (**Table 3**); small differences in one (243) were probably from threshold effects. This stability also showed in variant analysis, which demonstrated the absence of genotype switches in all donors. Such switches may be caused by the differential growth advantages of individual strains or intrahost recombinants, as reported recently in transplant recipients [22].

**Figure 2.**
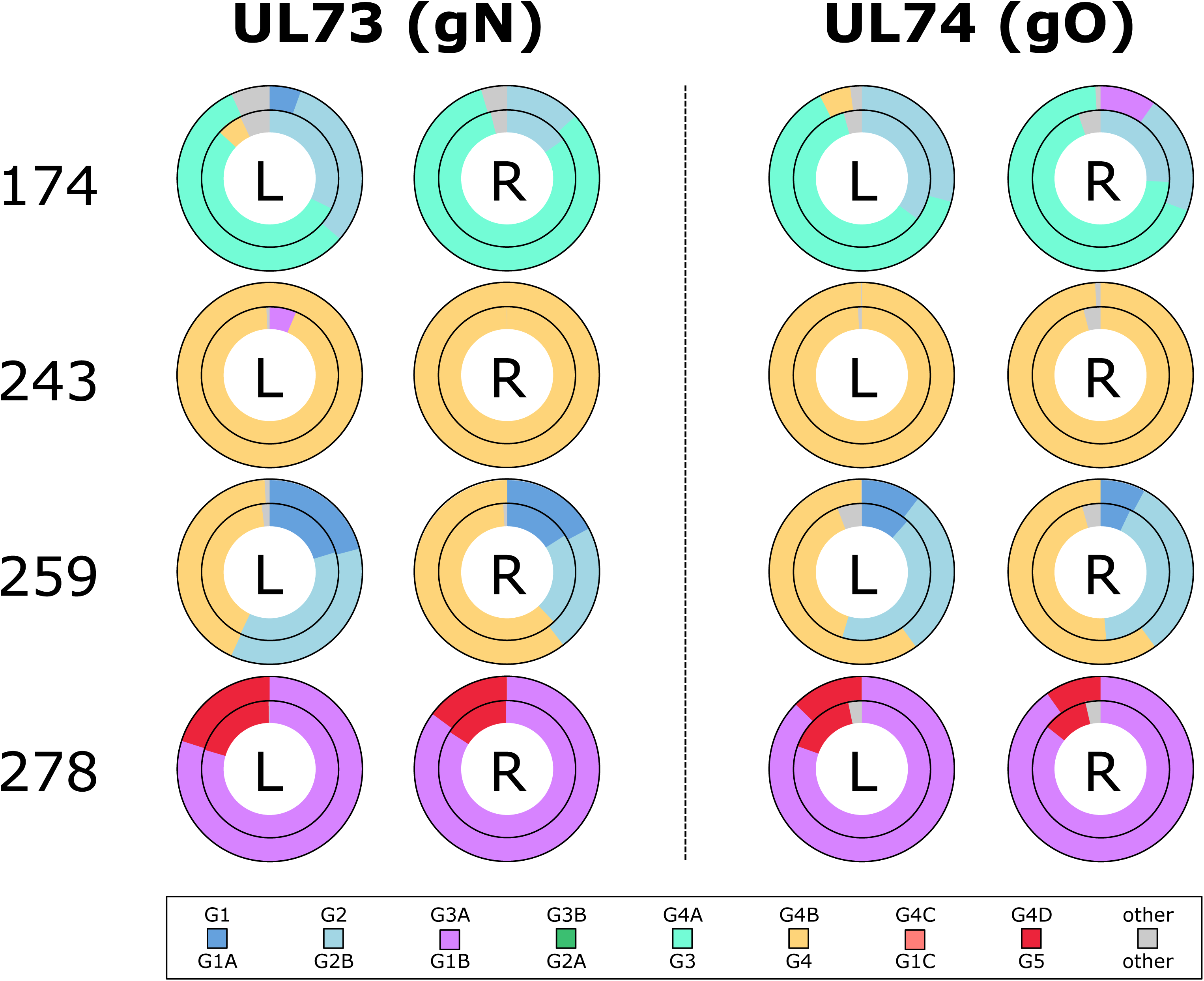
UL73 and UL74 genotypes in milk samples collected from the left (L) and right (R) breasts of four HIV-positive donors at 16 weeks postpartum (**Table 1**). The inner and outer rings show the results obtained using short and long motifs, respectively. Short motif 3’ was used for UL74 (**Table 2**). The colour key for genotypes is shown at the foot. Reads that did not meet the inclusion criteria for genotyping are shown as “other”.

Finally, additional long sequence motifs were used to investigate whether any strains contained gene-disrupting mutations detected previously in the set of 243 genomes [24]. These virus ‘pseudogene’ mutations cause frameshifting, resulting in loss or modification of the encoded proteins. They are more common in certain genes, most frequently and in order of decreasing abundance UL9, RL5A, UL1, RL6, US9 and UL111A [14, 16, 24]. The use of motifs representing three mutations in RL5A (present in 37 members of the 243 genome set), two in US9 (35 members) and one in UL111A (five members) demonstrated the presence of the RL5A and US9 mutations, but not that in UL111A, encoding viral interleukin-10 (**Supplementary Table 1** rows 167-174).

## DISCUSSION

Analysis of HCMV genomes directly from clinical samples is necessary for characterising infectious natural populations while avoiding the mutational artefacts arising from laboratory adaptation to cell-culture. Target enrichment has proven successful in this regard [11, 22, 24], but accurate genome analysis can be confounded by multiple strains, particularly in immunosuppressed groups in whom additional complexity may accumulate by reinfection or reactivation [17, 21, 24]. We have shown previously that HIV-positive women in sub-Saharan Africa have higher HCMV loads in breast milk than HIV-negative women, and that this is associated with adverse infant development [1, 3]. However, genomic studies of HCMV in milk samples or, indeed, samples from Africa, are scarce. We examined milk because of its importance in HCMV transmission, with the aim of understanding strain diversity and the burden of infection in HIV-positive (immunosuppressed) mothers, which may affect their infants. The sequence datasets were generated from 28 samples donated by 22 women, and 20 datasets from 14 women meeting quality thresholds were analysed.

The analysis focused on counting reads containing motifs specific to the genotypes of hypervariable genes. Short motifs were developed initially for sensitive characterisation of UL73 and UL74, which encode glycoproteins N and O (gN and gO), respectively, and then long motifs were used for further resolution of these two genes and ten others. Since UL73 and UL74 are linked and behave as a single genotype, as further shown here, haplotypes could not be determined using solely the short motifs (**Supplementary Table 2**) [7, 12]. However, mapping three short motifs to each UL74 genotype was uniquely useful for detecting intragenic recombination. The use of long motifs in a larger number of genes allowed increased resolution and haplotype information. These were less compromised by residual human reads in the datasets, but more susceptible to mismatches in target genomes (**Table 3**).

Genotypic and haplotypic complexity in this small cohort was remarkable. Most (82%) of the genotypes possible in the 12 hypervariable genes were detected, and 85% of the HIV-positive donors were infected by multiple strains. The level of multiple strain infection exceeds that in previous cohort analyses, including congenitally-infected and transplantation patients from developed countries, 243 set [22, 24]. Each of the eight fully characterised haplotypes identified was unique in this cohort and also in the set of 243 strains, in which most strains (223) are also unique [24]. In the milk cohort, up to 30 strains could be determined, using partial haplotypes. These observations testify to the huge number of HCMV haplotypes that may exist, possibly exceeding that related to immune diversity, as was recognised long before the high-throughput sequencing era [7, 8, 9, 12, 13, 15]. No evidence emerged for the existence of novel African genotypes, consistent with the view that HCMV genotypes are distributed throughout the world, although their relative prevalence may vary [7, 8, 12].

Strain composition in individual women was essentially stable, compartmentally (in milk samples from both breasts) and longitudinally (at 4 and 16 weeks postpartum). This indicates that the strains detected were present in the donor prior to viral reactivation in breast tissue during lactation. Leukocyte infiltrates have been characterised during this period [36], and may be the source of reactivated virus. A study conducted in Uganda has shown that mothers can be infected with strains from children [37]. However, even though multiple-strain infections were common in the cohort, there was limited evidence for reinfection or reactivation during the 4-16 week period postpartum. It is likely that opportunities for fresh infection existed at home and in the hospital, because all the mothers were HCMV-positive and had young children at home. The high proportion of multiple-strain infections observed also provides in principle for the diversification by recombination. These observations differ from those made in developed countries. A proportion of transplant-associated infections involve multiple strains, and these exhibit substantial longitudinal dynamism [22, 24] and are also associated with increased viral loads and the pathological outcomes of HCMV disease [18, 20]. The contrasting observation that most congenital or postnatal infections involve single strains [24, 38] suggests that only certain strains cross the placenta or are transmitted by breast milk, urine or saliva, perhaps due to the competence of a few virions to establish infection [38, 39]. This also implies that the HIV-positive women were exposed to a high burden of superinfection.

Whole-genome analyses and earlier PCR-based studies showed a high degree of linkage within the UL73/UL74 [7, 12, 24, 35] and RL12/RL13/UL1 loci [11, 24]. This is consistent with the involvement of homologous recombination during HCMV evolution, and may also reflect the functional constraints imposed on proteins that interact with each other or have interdependent functions. The UL73 and UL74 proteins (gN and gO) are part of the viral entry complex and have roles in viral exocytosis, cellular tropism and modulation of antibody neutralisation, and the RL12, RL13 and UL1 proteins are known or suspected to be involved in aspects of immune evasion probably mediated by an immunoglobulin-like binding domain shared by these proteins and other members of the RL11 family [9, 13, 40-43]. In addition, RL13 may influence the effect of UL74 on the growth of HCMV [44]. It is possible that different genotypes of hypervariable genes, and different combinations of genotypes, provide variable growth properties leading to higher viral loads and specific pathologies. For example, UL74 genotypes differentially affects viral growth properties *in vitro* [45], and genotypes of UL146, which is the most hypervariable gene in HCMV and encodes a vCXCL1 chemokine, affects neutrophil chemotaxis efficiency [46]. Similarly, host variation would be higher in Africa and may affect susceptibility as a result of HCMV genotype-specific interactions, for example with immunoglobulin variants [47-50].

Although data on genotypes and mutants could be extracted from the datasets regardless of strain complexity, complete genome sequence determination was possible for only four datasets because of high frequency of confounding multiple infections. To our knowledge, these are the first complete HCMV genome sequences to be determined from people living in Africa. Moreover, one of these originated from an HIV-negative woman and thus represents the first from an immunocompetent adult lacking HCMV-associated pathology. Future research is likely to focus on understanding the differences in HCMV transmission in immunosuppressed and immunocompetent settings in order to define the interplay between viral strain and host immunotype diversity in controlling disease. NOTES

## NOTES

### Acknowledgements

The authors thank the participants and clinical staff, managed by Dr L. Kasonka at the University Teaching Hospital, Lusaka, for facilitating the Breastfeeding and Postpartum Health Study, directed by Professor S. Filteau, London School of Hygiene and Tropical Medicine, for enabling the follow up analyses. We are grateful to Dr J. Hughes, MRC-University of Glasgow Centre for Virus Research, for advice on bioinformatic analysis, and Drs T. Clark and J. Phelan, London School of Hygiene and Tropical Medicine, for facilitating UNIX cluster access and Perl support.

### Supplementary data

Supplementary materials are available at The Journal of Infectious Diseases online. These consist of data provided by the authors to benefit the reader, are not copyedited, and are the sole responsibility of the authors. Questions or comments should be addressed to the corresponding author.

