## SupplementalTable3 for "Multiple-Strain Infections of Human Cytomegalovirus with High Genomic Diversity are Common In Breast Milk from HIV-Positive Women in Zambia"

| **Supplement Table 3.**  HCMV haplotypes of hypervariable genes are diverse between individuals and genotype prevalence shows restricted genetic linkage | | | | | | | | | | | | | | | |
| --- | --- | --- | --- | --- | --- | --- | --- | --- | --- | --- | --- | --- | --- | --- | --- |
|  | Genotypes hypervariable genes | | | | | | | | | | | | | | |
| ID | Mix | Strain | HIV | RL5A | RL6 | **RL12** | **RL13** | **UL1** | UL9 | UL11 | **UL73** | **UL74** | UL120 | UL146 | UL139 |
| 154 | major | z-a | - | **4** | **7** | **4A** | **4A** | **4** | **1** | **1** | **4A** | **3** | **4B** | **13** | **5** |
| 166 | single | z-b | - | **-** | **3** | **9** | **-** | **-** | **9** | **-** | **-** | **1A** | **4A** | **-** | **3** |
| 193 | single | z-c | - | **1** | **3** | **8** | **8** | **8** | **4** | **1** | **3A** | **1B** | **2B** | **12** | **3** |
| 232 | major | z-d | - | **1** | **1** | **1A** | **8** | **-** | **6** | **4** | **4D** | **5** | **4B** | **14** | **7** |
| 239 | major | z-e | - | **6** | **1** | **10** | **-** | **-** | **9** | **-** | **4B** | **-** | **2A** | **2** | **7** |
| 263 | single | z-f | - | **-** | **2** | **10** | **10** | **10** | **8** | **7** | **3A** | **1B** | **2A** | **-** | **4** |
| 280 | single | z-g | - | **1** | **-** | **-** | **6** | **6** | **7** | **-** | **1** | **1A** | **-** | **-** | **-** |
| 141 | major | z-h | + | **1** | **3** | **4B** | **2** | **4** | **2** | **6** | **2** | **2B** | **4B** | **12** | **3** |
| 141 | minor | z-i | + | **6** | **2** | **2** | **2** | **2** | **3** | **5** | **3A** | **1B** | **3B** | **2** | **8** |
| 173 | major | z-j | + | **1** | **3** | **8** | **8** | **8** | **4** | **1** | **3A** | **1B** | **1A** | **9** | **1A** |
| 173 | minor | z-k | + | **2** | **2** | **6** | **6** | **6** | **8** | **3** | **4A** | **3** | **4B** | **3** | **7** |
| 174 | major | z-l | + | **2** | **4** | **1B** | **1** | **1** | **4** | **1** | **4A** | **3** | **2B** | **9** | **5** |
| 174 | minor | z-m | + | **1** | **2** | **6** | **6** | **6** | **6** | **4** | **2** | **2B** | **3A** | **7** | **7** |
| 181 | major | z-n | + | **1** | **2** | **4A** | **6** | **6** | **6** | **1** | **4A** | **3** | **2A** | **8** | **4** |
| 243 | major | z-o | + | **1** | **6** | **2** | **2** | **2** | **3** | **6** | **4B** | **4** | **2A** | **8** | **7** |
| 248 | major | z-p | + | **1** | **1** | **4A** | **4A** | **4** | **1** | **1** | **3B** | **2A** | **4B** | **-** | **4** |
| 258 | major | z-q | + | **6** | **2** | **3** | **3** | **3** | **3** | **6** | **3A** | **1B** | **2B** | **9** | **2** |
| 259 | major | z-r | + | **1** | **3** | **8** | **8** | **8** | **3** | **1** | **4B** | **4** | **4B** | **9** | **8** |
| 259 | minor | z-s | + | **2** | **4** | **1A** | **1** | **1** | **1** | **6** | **2** | **2B** | **1A** | **12** | **2** |
| 264 | major | z-t | + | **1** | **2** | **10** | **10** | **10** | **8** | **7** | **4B** | **4** | **3A** | **10** | **3** |
| 264 | minor | z-u | + | **1** | **1** | **8** | **8** | **8** | **8** | **4** | **3A** | **1B** | **1A** | **10** | **5** |
| 277 | major | z-v | + | **1** | **3** | **4A** | **4A** | **4** | **9** | **1** | **4A** | **3** | **3A** | **1** | **3** |
| 277 | minor | z-w | + | **1** | **3** | **6** | **6** | **6** | **6** | **4** | **1** | **1A** | **4A** | **9** | **4** |
| 278 | major | z-x | + | **6** | **3** | **9** | **9** | **9** | **9** | **6** | **3A** | **1B** | **4B** | **8** | **2** |
| 278 | minor | z-y | + | **2** | **4** | **1A** | **1** | **1** | **1** | **1** | **4D** | **5** | **3A** | **9** | **2** |
| 281 | major | z-z | + | **6** | **3** | **1A** | **1** | **1** | **4** | **1** | **4D** | **5** | **4B** | **1** | **5** |
| 281 | minor | z-a1 | + | **1** | **3** | **6** | **6** | **6** | **7** | **3** | **3A** | **1B** | **3B** | **1** | **5** |
| 283 | major | z-b1 | + | **-** | **4** | **4B** | **4B** | **4** | **2** | **5** | **3A** | **1B** | **3A** | **1** | **5** |
| 288 | major | z-c1 | + | **2** | **4** | **6** | **6** | **6** | **4** | **1** | **4D** | **5** | **2B** | **3** | **4** |
| 288 | minor | z-d1 | + | **1** | **3** | **7** | **8** | **8** | **6** | **4** | **1** | **1A** | **4B** | **1** | **5** |
| Number of total genotypes | | | | 4/6 | 6/7 | 10/12 | 10/11 | 8/11 | 8/9 | 6/7 | 7/8 | 7/8 | 7/8 | 10/14 | 7/9 |
| Prevalence key | |  | >35% |  | >30% | >25% |  | >20% |  | >15% |  | >10% |  | >5% | >1% |
